# spatialLIBD: an R/Bioconductor package to visualize spatially-resolved transcriptomics data

**DOI:** 10.1101/2021.04.29.440149

**Authors:** Brenda Pardo, Abby Spangler, Lukas M. Weber, Stephanie C. Hicks, Andrew E. Jaffe, Keri Martinowich, Kristen R. Maynard, Leonardo Collado-Torres

## Abstract

**Motivation:** Spatially-resolved transcriptomics has now enabled the quantification of high-throughput and transcriptome-wide gene expression in intact tissue while also retaining the spatial coordinates. Incorporating the precise spatial mapping of gene activity advances our understanding of intact tissuespecific biological processes. In order to interpret these novel spatial data types, interactive visualization tools are necessary.

**Results:** We describe *spatialLIBD*, an R/Bioconductor package to interactively explore spatially-resolved transcriptomics data generated with the 10x Genomics Visium platform. The package contains functions to interactively access, visualize, and inspect the observed spatial gene expression data and data-driven clusters identified with supervised or unsupervised analyses, either on the user’s computer or through a web application.

**Availability:** *spatialLIBD* is available at bioconductor.org/packages/spatialLIBD.

**Supplementary information:** Supplementary data are available at *Bioinformatics* online.

## 1 Introduction

The rise of spatially-resolved transcriptomics technologies (Ståhl *et al.*, 2016; Xia *et al.*, 2019) are providing new opportunities to answer questions about the structure and function of complex intact tissues (Maynard *et al.*, 2020, 2021; Kuppe *et al.*, 2020). However, this exciting opportunity requires the development of new computational methods and interactive software. These tools have to accommodate the analysis of transcriptomic data with spatial coordinates and high resolution images into previously developed methods for single cell RNA-seq analysis (scRNA-seq), which lacked these components.

Currently there are limited options for interactive exploration of spatially-resolved transcriptomics data (Dries *et al.*, 2020). The 10x Genomics Loupe Browser (10x Genomics, 2020) utilizes a file created by the Space Ranger data pre-processing pipeline (10x Genomics, 2020) and performs marker gene analysis, as well as unsupervised graph-based and *k*-means clustering. Loupe visualizes both the spatially-resolved transcriptomics data along with the high resolution imaging data to export gene expression maps. An alternative to Loupe is the Giotto pipeline (Dries *et al.*, 2020), which contains both a data analysis module and a data visualization module. The data analysis module has many functions ranging from pre-processing to more advanced analysis such as characterizing cell-cell interactions. The visualization module contains an interactive workspace for exploring multiple layers of information. However, Loupe and Giotto currently do not support visualizing more than one tissue section at a time, which is useful for comparing replicates and annotating observed spots on the Visium array across samples. Furthermore, while unsupervised clustering methods are widely developed (Zhao *et al.*, 2020; Biancalani *et al.*, 2020), manual annotation of spots using known marker genes is important, as well as cross-sample dimension reduction techniques such as UMAP and t-SNE.

## 2 Results

*spatialLIBD* is an R/Bioconductor (Huber *et al.*, 2015) package that provides functions to interactively visualize and explore spatially-resolved transcriptomics data, including functionality to manually annotate spots. It was developed initially for visualizing multiple Visium tissue sections from the human brain (Maynard *et al.*, 2021), yet it is flexible enough to be used with other Visium data from different tissue types (Supplementary File 1, Figure 1A).

**Fig. 1.**
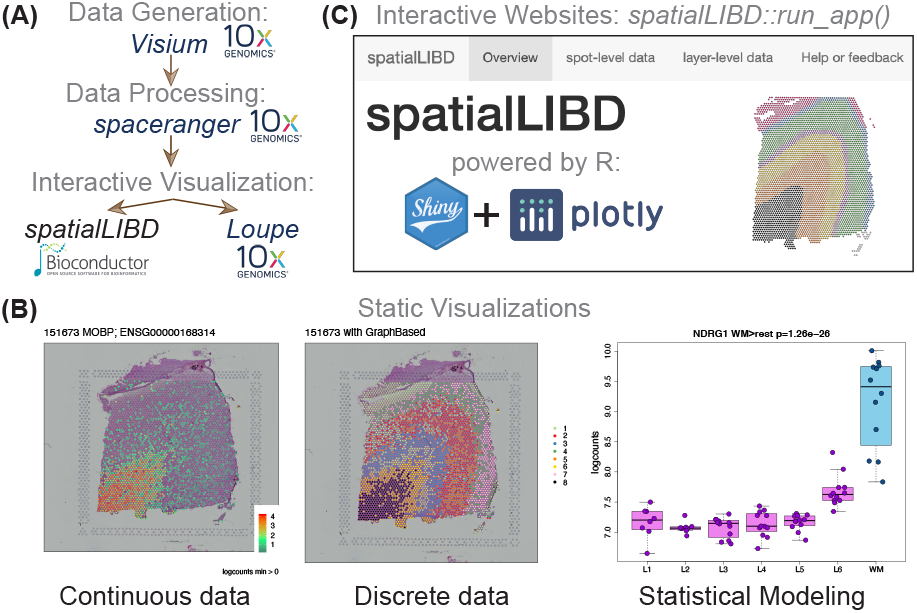
Overview of spatialLIBD: (A) Using spatially-resolved transcriptomics data generated with the 10x Visium platform and processed with spaceranger, both from 10x Genomics, spatialLIBD leverages the R/Bioconductor ecosystem and presents an alternative for interactive visualization to Loupe browser. (B) spatialLIBD supports static data visualizations of both continuous and categorical measurements along with the histology images, as well as visualizing results from downstream analyses. (C) spatialLIBD supports interactive websites for exploring the data powered by shiny and plotly.

The primary functions in *spatialLIBD* allow users (i) to visualize the spot-level spatial gene expression data, or any other continuous variable, and data-driven clusters or any categorical variable, with the histology image in the background (**Figure 1B**), and (ii) to inspect the data interactively, either on the user’s computer or through web hosting services such as shinyapps.io (**Figure 1C**). In addition, *spatialLIBD* enables users to visualize multiple samples at the same time, to export all static visualizations as PDF files or all interactive visualizations as PNG files, as well as all result tables as CSV files. Supplementary File 1 shows how to read in data processed with Space Ranger (10x Genomics, 2020) into R, make plots or create an interactive website with *spatialLIBD*.

Finally, *spatialLIBD* also supports the interactive exploration of data from Maynard *et al.* (2021). For example, using the customized website spatial.libd.org/spatialLIBD, researchers can perform spatial registration of their scRNA-seq clusters. This data can also be used for developing new analytical methods (Biancalani *et al.*, 2020; Zhao *et al.*, 2020).

## 3 Methods

*spatialLIBD* is designed to work with *SpatialExperiment* R/Bioconductor objects (Righelli *et al.*, 2021). It uses *shiny* (Chang *et al.*, 2021) and *plotly* (Sievert, 2020) for the interactive website, and *ggplot2* (Wickham, 2016) for the static images.

## 4 Discussion

*spatialLIBD* benefits from the open-source Bioconductor ecosystem (Huber *et al.*, 2015) and relies on the infrastructure provided by *SpatialExperiment* (Righelli *et al.*, 2021). It provides user-friendly functionality for visualizing continuous and categorical measurements along with the tissue images from spatially-resolved transcriptomics data from projects involving one or more samples measured on the 10x Genomics Visium platform. *spatialLIBD* provides a simple interactive website, which can be used for sharing data on the web, as well as manually annotating spots. *spatialLIBD* has limitations (Supplementary File 1) that are inherent to the methods used to implement it, such as (i) the memory per user required by a server for hosting the web application, (ii) response speeds for the interactive views due to the number of spots, (iii) the resolution of the images displayed limiting the usefulness to magnify specific spots, and (iv) customization of the web application by the end user. Despite these limitations, software solutions like *spatialLIBD* enable faster data exploration and insights in spatially-resolved transcriptomics research projects (Maynard *et al.*, 2021) and can serve as testing ground for ideas for future work, some of which are commercially available (BioTuring, 2021; 10x Genomics, 2020).

## Acknowledgements

Jesús Vélez Santiago and Ricardo O Ramírez-Flores from Julio Saez-Rodriguez’s group at Heidelberg University tested the applicability of *spatialLIBD* to new datasets and improve the interactive website code.

