## Supplementary File 1 for "spatialLIBD: an R/Bioconductor package to visualize spatially-resolved transcriptomics data"

### Using spatialLIBD with 10x Genomics public datasets

**29 April 2021**

**Package**

spatialLIBD 1.3.15

### 1 Basics

#### 1.1 Install `spatialLIBD`

`R` is an open-source statistical environment which can be easily modified to enhance its functionality via packages. *spatialLIBD* (Pardo, Spangler, Weber, Hicks, Jaffe, Martinowich, Maynard, and Collado-Torres, 2021) is an `R` package available via the Bioconductor repository for packages. `R` can be installed on any operating system from CRAN after which you can install *spatialLIBD* by using the following commands in your `R` session:

```
if (!requireNamespace("BiocManager", quietly = TRUE)) {
    install.packages("BiocManager")
}

BiocManager::install("spatialLIBD")

## Check that you have a valid Bioconductor installation
BiocManager::valid()
```

#### 1.2 Required knowledge

Please first check the *Introduction to spatialLIBD* vignette available through GitHub or Bioconductor.

#### 1.3 Citing `spatialLIBD`

We hope that *spatialLIBD* will be useful for your research. Please use the following information to cite the package and the overall approach. Thank you!

```
## Citation info
citation("spatialLIBD")
#>
#> Pardo B, Spangler A, Weber LM, Hicks SC, Jaffe AE, Martinowich K,
#> Maynard KR, Collado-Torres L (2021). _spatialLIBD: an R/Bioconductor
#> package to visualize spatially-resolved transcriptomics data_. doi:
#> 10.18129/B9.bioc.spatialLIBD (URL:
#> https://doi.org/10.18129/B9.bioc.spatialLIBD),
#> https://github.com/LieberInstitute/spatialLIBD - R package version
#> 1.3.15, <URL: http://www.bioconductor.org/packages/spatialLIBD>.
#>
#> Maynard KR, Collado-Torres L, Weber LM, Uytingco C, Barry BK, Williams
#> SR, II JLC, Tran MN, Besich Z, Tippani M, Chew J, Yin Y, Kleinman JE,
#> Hyde TM, Rao N, Hicks SC, Martinowich K, Jaffe AE (2021).
#> "Transcriptome-scale spatial gene expression in the human dorsolateral
#> prefrontal cortex." _Nature Neuroscience_. doi:
#> 10.1038/s41593-020-00787-0 (URL:
#> https://doi.org/10.1038/s41593-020-00787-0), <URL:
#> https://www.nature.com/articles/s41593-020-00787-0>.
#>
#> To see these entries in BibTeX format, use 'print(<citation>,
```

```
#> bibtex=TRUE)', 'toBibtex(.)', or set
#> 'options(citation.bibtex.max=999)'.
```

### 2 Download data from 10x Genomics

In this vignette we'll show you how you can use *spatialLIBD* (Pardo, Spangler, Weber, et al., 2021) for exploring spatially resolved transcriptomics data from the Visium platform by 10x Genomics. That is, you will learn how to use *spatialLIBD* for data beyond the one it was initially developed for (Maynard, Collado-Torres, Weber, Uytingco, Barry, Williams, II, Tran, Besich, Tippani, Chew, Yin, Kleinman, Hyde, Rao, Hicks, Martinowich, and Jaffe, 2021). To illustrate these steps, we will use data that 10x Genomics made publicly available at https://support.10xgenomics.com/spatial-gene-expression/datasets. We will use files from the human lymph node example publicly available at https://support.10xgenomics.com/spatial-gene-expression/datasets/1.1.0/V1_Human_Lymph_Node.

#### 2.1 Load packages

To get started, lets load the different packages we'll need for this vignette. Here's a brief summary of why we need these packages:

- *BiocFileCache*: for downloading and saving a local cache of the data
- *SpatialExperiment*: for reading the `spaceranger` files provided by 10x Genomics
- *rtracklayer*: for importing a gene annotation GTF file
- *pryr*: for checking how much memory our object is using
- *spatialLIBD*: for launching an interactive website to explore the data

```r
## Load packages in the order we'll use them next
library("BiocFileCache")
library("SpatialExperiment")
library("rtracklayer")
library("pryr")
library("spatialLIBD")
```

#### 2.2 Download spaceranger output files

Next we download data from 10x Genomics available from the human lymph node example, available at https://support.10xgenomics.com/spatial-gene-expression/datasets/1.1.0/V1_Human_Lymph_Node. We don't need to download all the files listed there since *SpatialExperiment* doesn't need all of them for importing the data into R. These files are part of the output that gets generated by `spaceranger` which is the processing pipeline provided by 10x Genomics for Visium data.

We'll use *BiocFileCache* to keep the data in a local cache in case we want to run this example again and don't want to re-download the data from the web.

```r
## Download and save a local cache of the data provided by 10x Genomics
bfc <- BiocFileCache::BiocFileCache()
lymph.url <-
    paste0(
        "https://cf.10xgenomics.com/samples/spatial-exp/",
```

```
        "1.1.0/V1_Human_Lymph_Node/",
        c(
            "V1_Human_Lymph_Node_filtered_feature_bc_matrix.tar.gz",
            "V1_Human_Lymph_Node_spatial.tar.gz"
        )
    )
lymph.data <- sapply(lymph.url, BiocFileCache::bfcrpath, x = bfc)
```

10x Genomics provides the files in compressed tarballs (`.tar.gz` file extension). Which is why we'll need to use `utils::untar()` to decompress the files. This will create new directories and we will use `list.files()` to see what files these directories contain.

```
## Extract the files to a temporary location
## (they'll be deleted once you close your R session)
sapply(lymph.data, utils::untar, exdir = tempdir())
#> https://cf.10xgenomics.com/samples/spatial-exp/1.1.0/V1_Human_Lymph_Node/V1_Human_Lymph_Node_filtered_fea
#>
#>                     https://cf.10xgenomics.com/samples/spatial-exp/1.1.0/V1_Human_Lymph_Node/V1_Human_Lymph
#>

## List the files we downloaded and extracted
## These files are typically SpaceRanger outputs
lymph.dirs <- file.path(
    tempdir(),
    c("filtered_feature_bc_matrix", "spatial", "raw_feature_bc_matrix")
)
list.files(lymph.dirs)
#> [1] "aligned_fiducials.jpg"     "barcodes.tsv.gz"
#> [3] "detected_tissue_image.jpg" "features.tsv.gz"
#> [5] "matrix.mtx.gz"             "scalefactors_json.json"
#> [7] "tissue_hires_image.png"    "tissue_lowres_image.png"
#> [9] "tissue_positions_list.csv"
```

Now that we have the files that we need, we can import the data into R using `read10xVisium()` from *SpatialExperiment*. We'll import the low resolution histology images produced by `spaceranger` using the `images = "lowres"` and `load = TRUE` arguments. We'll also load the filtered gene expression data using the `data = "filtered"` argument. The count matrix can still be quite large, which is why we'll use the `type = "sparse"` argument to load the data into an R object that is memory-efficient for sparse data.

```
## Import the data as a SpatialExperiment object
spe <- SpatialExperiment::read10xVisium(
    samples = tempdir(),
    sample_id = "lymph",
    type = "sparse", data = "filtered",
    images = "lowres", load = TRUE
)
## Inspect the R object we just created: class, memory, and how it looks in
## general
class(spe)
#> [1] "SpatialExperiment"
```

```
#> attr(,"package")
#> [1] "SpatialExperiment"
pryr::object_size(spe)
#> 296 MB
spe
#> class: SpatialExperiment
#> dim: 36601 4035
#> metadata(0):
#> assays(1): counts
#> rownames(36601): ENSG00000243485 ENSG00000237613 ... ENSG00000278817
#>   ENSG00000277196
#> rowData names(1): symbol
#> colnames(4035): AAACAAGTATCTCCCA-1 AAACAATCTACTAGCA-1 ...
#>   TTGTTTGTATTACACG-1 TTGTTTGTGTAAATTC-1
#> colData names(3): Sample Barcode sample_id
#> reducedDimNames(0):
#> mainExpName: NULL
#> altExpNames(0):
#> spatialData names(5) : in_tissue array_row array_col pxl_col_in_fullres
#>   pxl_row_in_fullres
#> imgData names(4): sample_id image_id data scaleFactor

## The counts are saved in a sparse matrix R object
class(counts(spe))
#> [1] "dgCMatrix"
#> attr(,"package")
#> [1] "Matrix"
```

### 3 Modify spe for spatialLIBD

Now that we have an `SpatialExperiment` R object (`spe`) with the data from 10x Genomics for the human lymph node example, we need to add a few features to the R object in order to create the interactive website using `spatialLIBD::run_app()`. These additional elements power features in the interactive website that you might be interested in.

First we start with adding a few variables to the sample information table (`colData()`) of our `spe` object. We add:

- `key`: this labels each spot with a unique identifier. We combine the sample ID with the spot barcode ID to create this unique identifier.
- `sum_umi`: this continuous variable contains the total number of counts for each sample prior to filtering any genes.
- `sum_gene`: this continuous variable contains the number of genes that have at least 1 count.

```
## Add some information used by spatialLIBD
spe$key <- paste0(spe$sample_id, "_", spe$Barcode)
spe$sum_umi <- colSums(counts(spe))
spe$sum_gene <- colSums(counts(spe) > 0)
```

#### 3.1 Add gene annotation information

The files `SpatialExperiment::read10xVisium()` uses to read in the `spaceranger` outputs into R do not include much information about the genes, such as their chromosomes, coordinates, and other gene annotation information. We thus recommend that you read in this information from a gene annotation file: typically a `gtf` file. For a real case scenario, you'll mostly likely have access to the GTF file provided by 10x Genomics. However, we cannot download that file without downloading other files for this example. Thus we'll show you the code you would use if you had access to the GTF file from 10x Genomics and also show a second approach that works for this vignette.

##### 3.1.1 From 10x

Depending on the version of `spaceranger` you used, you might have used different GTF files 10x Genomics has made available at https://support.10xgenomics.com/single-cell-gene-expression/software/downloads/latest and described at https://support.10xgenomics.com/single-cell-gene-expression/software/release-notes/build. These files are too big though and we won't download them in this example. For instance, *References - 2020-A (July 7, 2020)* for *Human reference (GRCh38)* is 11 GB in size and contains files we do not need for this vignette. If you did have the file locally, you could use the following code to read in the GTF file prepared by 10x Genomics and add the information into your `spe` object that `SpatialExperiment::read10xVisium()` does not include.

For example, in our computing cluster this GTF file is located at the following path and is 1.4 GB in size:

```
$ cd /dcl02/lieber/ajaffe/SpatialTranscriptomics/
$ du -sh --apparent-size refdata-gex-GRCh38-2020-A/genes/genes.gtf
1.4G    /dcl02/lieber/ajaffe/SpatialTranscriptomics/refdata-gex-GRCh38-2020-A/genes/genes.gtf
```

If you have the GTF file from 10x Genomics, we show next how you can read the information into R, match it appropriately with the information in the `spe` object and add it back into the `spe` object.

```
## You could:
## * download the 11 GB file from
##https://cf.10xgenomics.com/supp/cell-exp/refdata-gex-GRCh38-2020-A.tar.gz
## * decompress it

## Read in the gene information from the annotation GTF file provided by 10x
gtf <-
    rtracklayer::import(
        "/path/to/refdata-gex-GRCh38-2020-A/genes/genes.gtf"
    )

## Subject to genes only
gtf <- gtf[gtf$type == "gene"]

## Set the names to be the gene IDs
names(gtf) <- gtf$gene_id

## Match the genes
match_genes <- match(rownames(spe), gtf$gene_id)
```

```r
## They should all be present if you are using the correct GTF file from 10x
stopifnot(all(!is.na(match_genes)))

## Keep only some columns from the gtf (you could keep all of them if you want)
mcols(gtf) <-
    mcols(gtf)[, c(
        "source",
        "type",
        "gene_id",
        "gene_version",
        "gene_name",
        "gene_type"
    )]

## Add the gene info to our SPE object
rowRanges(spe) <- gtf[match_genes]

## Inspect the gene annotation data we added
rowRanges(spe)
```

##### 3.1.2 From Gencode

In this vignette, we'll use the GTF file from Gencode v32. That's because the build notes from *References - 2020-A (July 7, 2020)* and *Human reference, GRCh38 (GENCODE v32/Ensembl 98)* at https://support.10xgenomics.com/single-cell-gene-expression/software/release-notes/build#GRCh38_2020A show that 10x Genomics used Gencode v32. They also used Ensembl version 98 which is why a few genes we have in our object are going to be missing. We show next how you can read the information into R, match it appropriately with the information in the `spe` object and add it back into the `spe` object.

```r
## Download the Gencode v32 GTF file and cache it
gtf_cache <- BiocFileCache::bfcrpath(
    bfc,
    paste0(
        "ftp://ftp.ebi.ac.uk/pub/databases/gencode/Gencode_human/",
        "release_32/gencode.v32.annotation.gtf.gz"
    )
)

## Show the GTF cache location
gtf_cache
#>                                                                                                   BF[illegible]
#> "/Users/lcollado/Library/Caches/org.R-project.R/R/BiocFileCache/665c5ccf84d7_gencode.v32.annotation.gtf.g[illegible]

## Import into R (takes ~1 min)
gtf <- rtracklayer::import(gtf_cache)

## Subset to genes only
gtf <- gtf[gtf$type == "gene"]
```

```
## Remove the .x part of the gene IDs
gtf$gene_id <- gsub("\\..*", "", gtf$gene_id)

## Set the names to be the gene IDs
names(gtf) <- gtf$gene_id

## Match the genes
match_genes <- match(rownames(spe), gtf$gene_id)
table(is.na(match_genes))
#>
#> FALSE  TRUE
#> 36572    29

## Drop the few genes for which we don’t have information
spe <- spe[!is.na(match_genes), ]
match_genes <- match_genes[!is.na(match_genes)]

## Keep only some columns from the gtf
mcols(gtf) <- mcols(gtf)[, c("source", "type", "gene_id", "gene_name", "gene_type")]

## Add the gene info to our SPE object
rowRanges(spe) <- gtf[match_genes]

## Inspect the gene annotation data we added
rowRanges(spe)
#> GRanges object with 36572 ranges and 5 metadata columns:
#>                   seqnames        ranges strand |   source     type
#>                      <Rle>     <IRanges>  <Rle> | <factor> <factor>
#>   ENSG00000243485     chr1   29554-31109      + |   HAVANA     gene
#>   ENSG00000237613     chr1   34554-36081      - |   HAVANA     gene
#>   ENSG00000186092     chr1   65419-71585      + |   HAVANA     gene
#>   ENSG00000238009     chr1 89295-133723      - |   HAVANA     gene
#>   ENSG00000239945     chr1   89551-91105      - |   HAVANA     gene
#>               ...      ...           ...    ... .      ...      ...
#>   ENSG00000212907     chrM   10470-10766      + |  ENSEMBL     gene
#>   ENSG00000198886     chrM   10760-12137      + |  ENSEMBL     gene
#>   ENSG00000198786     chrM   12337-14148      + |  ENSEMBL     gene
#>   ENSG00000198695     chrM   14149-14673      - |  ENSEMBL     gene
#>   ENSG00000198727     chrM   14747-15887      + |  ENSEMBL     gene
#>                           gene_id   gene_name      gene_type
#>                       <character> <character>    <character>
#>   ENSG00000243485 ENSG00000243485 MIR1302-2HG         lncRNA
#>   ENSG00000237613 ENSG00000237613     FAM138A         lncRNA
#>   ENSG00000186092 ENSG00000186092       OR4F5 protein_coding
#>   ENSG00000238009 ENSG00000238009  AL627309.1         lncRNA
#>   ENSG00000239945 ENSG00000239945  AL627309.3         lncRNA
#>               ...             ...         ...            ...
#>   ENSG00000212907 ENSG00000212907     MT-ND4L protein_coding
#>   ENSG00000198886 ENSG00000198886      MT-ND4 protein_coding
#>   ENSG00000198786 ENSG00000198786      MT-ND5 protein_coding
#>   ENSG00000198695 ENSG00000198695      MT-ND6 protein_coding
```

```
#>   ENSG00000198727 ENSG00000198727      MT-CYB protein_coding
#>   -------
#>   seqinfo: 25 sequences from an unspecified genome; no seqlengths
```

##### 3.1.3 Enable a friendlier gene search

Regardless of which method you used to obtain the gene annotation information, we can now proceed by adding the gene symbol and gene ID information that helps users search for genes in the shiny app produced by `spatialLIBD`. This will enable users to search genes by gene symbol or gene ID. If you didn't do this, users would only be able to search genes by gene ID which makes the web application harder to use.

We also compute the total expression for the mitochondrial chromosome (chrM) as well as the ratio of chrM expression. Both of these continuous variables are interesting to explore and in some situations could be useful for biological interpretations. For instance, in our pilot data (Maynard, Collado-Torres, Weber, et al., 2021), we noticed that the `expr_chrM_ratio` was associated to DLPFC layers. That is, spots with high `expr_chrM_ratio` were not randomly located in our Visium slides.

```
## Add information used by spatialLIBD
rowData(spe)$gene_search <- paste0(
    rowData(spe)$gene_name, "; ", rowData(spe)$gene_id
)

## Compute chrM expression and chrM expression ratio
is_mito <- which(seqnames(spe) == "chrM")
spe$expr_chrM <- colSums(counts(spe)[is_mito, , drop = FALSE])
spe$expr_chrM_ratio <- spe$expr_chrM / spe$sum_umi
```

#### 3.2 Filter the spe object

We can now continue with some filtering steps since this can help reduce the object size in memory as well as make it ready to use for downstream processing tools such as those from the *scran* and *scuttle* packages. Though these steps are not absolutely necessary.

```
## Remove genes with no data
no_expr <- which(rowSums(counts(spe)) == 0)

## Number of genes with no counts
length(no_expr)
#> [1] 11397

## Compute the percent of genes with no counts
length(no_expr) / nrow(spe) * 100
#> [1] 31.16318
spe <- spe[-no_expr, , drop = FALSE]

## Remove spots without counts
summary(spe$sum_umi)
#>    Min. 1st Qu.  Median    Mean 3rd Qu.    Max.
#>      23   15917   20239   20738   25252   54931
```

```
## If we had spots with no counts, we would remove them
if (any(spe$sum_umi == 0)) {
    spots_no_counts <- which(spe$sum_umi == 0)
    ## Number of spots with no counts
    print(length(spots_no_counts))
    ## Percent of spots with no counts
    print(length(spots_no_counts) / ncol(spe) * 100)
    spe <- spe[, -spots_no_counts, drop = FALSE]
}
```

#### 3.3 Check object

Next, we add the `ManualAnnotation` variable to the sample information table (`colData()`) with `"NA"`. That variable is used by the interactive website to store any manual annotations.

```
## Add a variable for saving the manual annotations
spe$ManualAnnotation <- "NA"
```

Finally, we can now check the final object using `spatialLIBD::check_spe()`. This is a helper function that will warn us if some important element is missing in `spe` that we use later for the interactive website. If it all goes well, it will return the original `spe` object.

```
## Check the final dimensions and object size
dim(spe)
#> [1] 25175  4035
pryr::object_size(spe)
#> 298 MB

## Run check_spe() function
spatialLIBD::check_spe(spe)
#> class: SpatialExperiment
#> dim: 25175 4035
#> metadata(0):
#> assays(1): counts
#> rownames(25175): ENSG00000238009 ENSG00000241860 ... ENSG00000198695
#>   ENSG00000198727
#> rowData names(6): source type ... gene_type gene_search
#> colnames(4035): AAACAAGTATCTCCCA-1 AAACAATCTACTAGCA-1 ...
#>   TTGTTTGTATTACACG-1 TTGTTTGTGTAAATTC-1
#> colData names(9): Sample Barcode ... expr_chrM_ratio ManualAnnotation
#> reducedDimNames(0):
#> mainExpName: NULL
#> altExpNames(0):
#> spatialData names(5) : in_tissue array_row array_col pxl_col_in_fullres
#>   pxl_row_in_fullres
#> imgData names(4): sample_id image_id data scaleFactor
```

### 4 Explore the data

With our complete `spe` object, we can now use *spatialLIBD* for visualizing our data. We can do so using functions such as `vis_gene()` and `vis_clus()` that are described in more detail at the *Introduction to spatialLIBD* vignette available through GitHub or Bioconductor.

```
# Visualizations
spatialLIBD::vis_gene(
    spe = spe,
    sampleid = "lymph",
    geneid = "sum_umi",
    assayname = "counts"
)
```

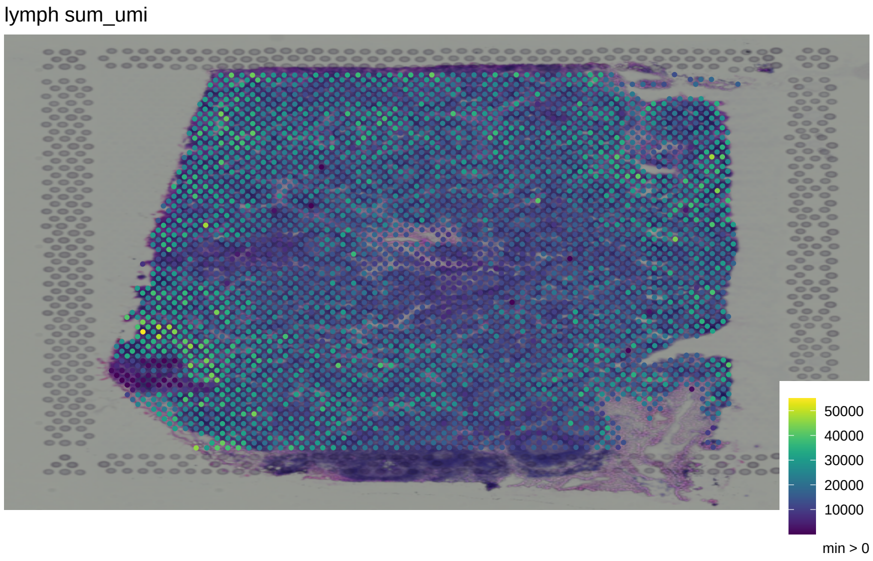

```
set.seed(20210428)
spe$random_cluster <- sample(1:7, ncol(spe), replace = TRUE)
spatialLIBD::vis_clus(
    spe = spe,
    sampleid = "lymph",
    clustervar = "random_cluster"
)
```

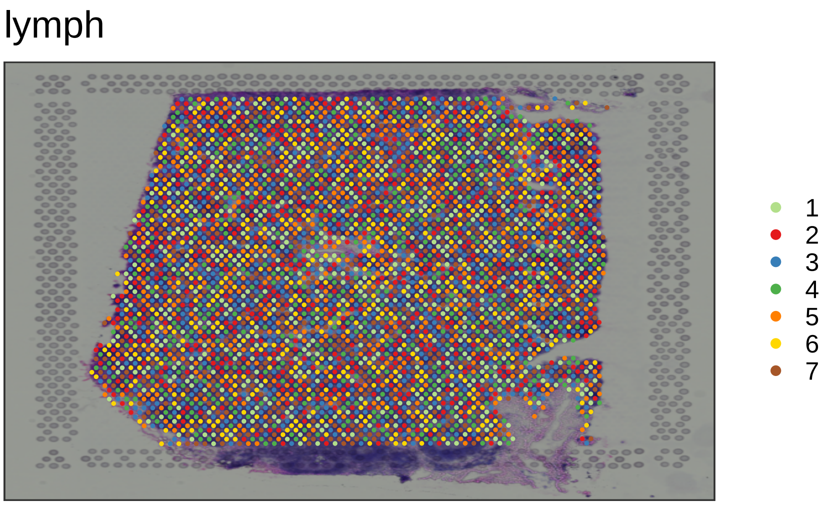

#### 4.1 Run the interactive website

We are now ready to create our interactive website for the human lymph node data. The interactive website is a *shiny* web application that uses *plotly* to power several of the interactive features. We can create the interactive website using the `spatialLIBD::run_app()` function. The default arguments of that function are customized for the data from our initial study (Maynard, Collado-Torres, Weber, et al., 2021), so we will need to make some adjustments:

- `sce_layer`, `modeling_results` and `sig_genes` will be set to `NULL` since do not have any pseudo-bulk results for this example data.
- `title`: we will use a custom title that reflect our data
- `spe_discreate_vars`: we don't have really any discrete variables to show beyond `ManualAnnotation` which is used for the manual annotations and `random_cluster` that we created in the previous section.
- `spe_continous_vars`: we have computed several continuous variables while adapting our `spe` object for *spatialLIBD*, so we'll list these variables below in order to visually inspect them.
- `default_cluster`: this is used for indicating the default discrete variable and for now we'll set it to our `random_cluster`.

```
## Run our shiny app
if (interactive()) {
    run_app(
        spe,
        sce_layer = NULL,
        modeling_results = NULL,
        sig_genes = NULL,
        title = "spatialLIBD: human lymph node by 10x Genomics",
        spe_discrete_vars = c("random_cluster", "ManualAnnotation"),
        spe_continuous_vars = c("sum_umi", "sum_gene", "expr_chrM", "expr_chrM_ratio"),
        default_cluster = "random_cluster"
    )
}
```

We also recommend creating custom website documentation files as described in the documentation of `spatialLIBD::run_app()`. Those documentation files will help you describe your project to your users in a more personalized way. The easiest way to start is to copy our documentation files to a new location and adapt them. You can located them at the following path.

```
## Locate our documentation files
docs_path <- system.file("app", "www", package = "spatialLIBD")
docs_path
#> [1] "/Library/Frameworks/R.framework/Versions/4.1/Resources/library/spatialLIBD/app/www"
list.files(docs_path)
#> [1] "documentation_sce_layer.md" "documentation_spe.md"
#> [3] "favicon.ico"                "footer.html"
#> [5] "README.md"
```

### 5 Limitations

`spatialLIBD::run_app()` has limitations that are inherent to the methods used to implement it, such as:

1. the memory per user required by a server for hosting the web application,
2. response speeds for the interactive views due to the number of spots,
3. the resolution of the images displayed limiting the usefulness to magnify specific spots,
4. and customization of the web application by the end user.

#### 5.1 Memory

Regarding the memory limitation, you can estimate how much memory you need per user by considering the memory required for the `spe` and `sce_layer` objects.

```
pryr::object_size(spe)
#> 298 MB
```

In our pilot data (Maynard, Collado-Torres, Weber, et al., 2021) our object uses about 2.1 GB of RAM since it contains the data for 12 Visium slides and we considered using about 3 GB of RAM per user. You could filter the genes more aggressively to drop lowly expressed genes or if you have many Visium slides, you could consider making multiple websites for different sets of slides. You could also have multiple mirrors to support several users, though in that case, we recommend linking users to a stable website instead of one that might not be available if you have too many users: for us our stable website is http://research.libd.org/spatialLIBD/ which includes the links to all the mirrors.

Given these memory limitations, we chose to deploy our main web application at http://spatial.libd.org/spatialLIBD/ using an Amazon EC2 instance: an 'r5.4xlarge' EC2 instance with 16 vCPUs, 128 GB DRAM, 10 Gb max network, 1.008 USD/Hour. We also have deployed mirrors at https://www.shinyapps.io/ using the "3X-Large (8 GB)" instances they provide.

#### 5.2 Response speeds

This limitation is mostly due to the number of spots shown under the "clusters (interactive)" section of the interactive website powered by *plotly*. Each spot is shown four times which is about 16 thousand spots for one Visium slide (depending on any filter steps you applied). The response time will depend on your own computer RAM memory, that is, the *client* side. This limitation might be more noticeable if you have a computer with 8GB of RAM or lower, as well as if you have other high-memory software open. Furthermore, if you are running web application locally through `spatialLIBD::run_app()` then you also need to consider the required memory for the R objects. That is, the *server* side memory use.

Thanks to Jesús Vélez Santiago, the app is more responsive as of version 1.3.15 by using `plotly::toWebGL()`.

#### 5.3 Image resolution

When you construct the SpatialExperiment `spe` object with *SpatialExperiment*, you can read in higher resolution images. However, the benefit of loading the raw histology images (500 MB to 20 GB per image) is likely non-existent in this web application. The memory required would likely become prohibitive. Other solutions load these raw histology images in chunks and display the chunks necessary for a given visualization area. We thus recommend using other software if you want to zoom in at the spot and/or cell resolution.

#### 5.4 Customization

While the documentation, title, icon and HTML footer are all customizable at `spatialLIBD::run_app()`, ultimately the panels shown are not unless you fork and adapt the internal code of this package. Thus, the interactive web applications powered by *spatialLIBD* are not as easy to customize as say *iSEE* web applications are. We think of our web application as a good enough prototype that can be useful for initial explorations of 10x Genomics Visium data. We welcome additions to our code, though we recognize that you might want to build your own production-level solution.

### 6 Reproducibility

The *spatialLIBD* package (Pardo, Spangler, Weber, et al., 2021) was made possible thanks to:

- R (R Core Team, 2021)
- *BiocFileCache* (Shepherd and Morgan, 2021)
- *BiocStyle* (Oleś, 2021)
- *knitr* (Xie, 2021)
- *pryr* (Wickham, 2018)
- *RefManageR* (McLean, 2017)
- *rmarkdown* (Allaire, Xie, McPherson, Luraschi, Ushey, Atkins, Wickham, Cheng, Chang, and Iannone, 2021)
- *rtracklayer* (Lawrence, Gentleman, and Carey, 2009)
- *sessioninfo* (Csárdi, core, Wickham, Chang, Flight, Müller, and Hester, 2018)
- *SpatialExperiment* (Righelli and Risso, 2021)
- *testthat* (Wickham, 2011)

This package was developed using *biocthis*.

Code for creating the vignette

```
## Create the vignette
library("rmarkdown")
system.time(render("TenX_data_download.Rmd", "BiocStyle::html_document"))

## Extract the R code
library("knitr")
knit("TenX_data_download.Rmd", tangle = TRUE)
```

Date the vignette was generated.

```
#> [1] "2021-04-29 14:50:26 EDT"
```

Wallclock time spent generating the vignette.

```
#> Time difference of 2.579 mins
```

R session information.

```
#> - Session info -----------------------------------------------------------------------------------------------
#>  setting  value
#>  version  R version 4.1.0 alpha (2021-04-26 r80229)
#>  os       macOS Big Sur 10.16
#>  system   x86_64, darwin17.0
#>  ui       X11
#>  language (EN)
#>  collate  en_US.UTF-8
#>  ctype    en_US.UTF-8
#>  tz       America/New_York
#>  date     2021-04-29
#>
#> - Packages ---------------------------------------------------------------------------------------------------
#>  package              * version    date       lib source
#>  AnnotationDbi          1.53.1     2021-02-04 [1] Bioconductor
#>  AnnotationHub          2.99.5     2021-04-16 [1] Bioconductor
#>  assertthat             0.2.1      2019-03-21 [1] CRAN (R 4.1.0)
#>  attempt                0.3.1      2020-05-03 [1] CRAN (R 4.1.0)
#>  beachmat               2.7.7      2021-02-22 [1] Bioconductor
#>  beeswarm               0.3.1      2021-03-07 [1] CRAN (R 4.1.0)
#>  benchmarkme            1.0.7      2021-03-21 [1] CRAN (R 4.1.0)
#>  benchmarkmeData        1.0.4      2020-04-23 [1] CRAN (R 4.1.0)
#>  Biobase              * 2.51.0     2020-10-27 [1] Bioconductor
#>  BiocFileCache        * 1.99.6     2021-04-16 [1] Bioconductor
#>  BiocGenerics         * 0.37.4     2021-04-23 [1] Bioconductor
#>  BiocIO                 1.1.2      2020-12-05 [1] Bioconductor
#>  BiocManager            1.30.12    2021-03-28 [1] CRAN (R 4.1.0)
#>  BiocNeighbors          1.9.4      2020-12-17 [1] Bioconductor
#>  BiocParallel           1.25.5     2021-03-09 [1] Bioconductor
#>  BiocSingular           1.7.2      2021-01-23 [1] Bioconductor
#>  BiocStyle            * 2.19.2     2021-03-28 [1] Bioconductor
```

```
#>  BiocVersion            3.13.1    2020-10-27 [1] Bioconductor
#>  Biostrings             2.59.2    2020-12-18 [1] Bioconductor
#>  bit                    4.0.4     2020-08-04 [1] CRAN (R 4.1.0)
#>  bit64                  4.0.5     2020-08-30 [1] CRAN (R 4.1.0)
#>  bitops                 1.0-7     2021-04-24 [1] CRAN (R 4.1.0)
#>  blob                   1.2.1     2020-01-20 [1] CRAN (R 4.1.0)
#>  bookdown               0.22      2021-04-22 [1] CRAN (R 4.1.0)
#>  bslib                  0.2.4     2021-01-25 [1] CRAN (R 4.1.0)
#>  cachem                 1.0.4     2021-02-13 [1] CRAN (R 4.1.0)
#>  cli                    2.5.0     2021-04-26 [1] CRAN (R 4.1.0)
#>  codetools              0.2-18    2020-11-04 [1] CRAN (R 4.1.0)
#>  colorout               1.2-2     2020-11-03 [1] Github (jalvesaq/colorout@726d681)
#>  colorspace             2.0-0     2020-11-11 [1] CRAN (R 4.1.0)
#>  config                 0.3.1     2020-12-17 [1] CRAN (R 4.1.0)
#>  cowplot                1.1.1     2020-12-30 [1] CRAN (R 4.1.0)
#>  crayon                 1.4.1     2021-02-08 [1] CRAN (R 4.1.0)
#>  curl                   4.3       2019-12-02 [1] CRAN (R 4.1.0)
#>  data.table             1.14.0    2021-02-21 [1] CRAN (R 4.1.0)
#>  DBI                    1.1.1     2021-01-15 [1] CRAN (R 4.1.0)
#>  dbplyr               * 2.1.1     2021-04-06 [1] CRAN (R 4.1.0)
#>  DelayedArray           0.17.10   2021-03-24 [1] Bioconductor
#>  DelayedMatrixStats     1.13.6    2021-04-22 [1] Bioconductor
#>  desc                   1.3.0     2021-03-05 [1] CRAN (R 4.1.0)
#>  digest                 0.6.27    2020-10-24 [1] CRAN (R 4.1.0)
#>  dockerfiler            0.1.3     2019-03-19 [1] CRAN (R 4.1.0)
#>  doParallel             1.0.16    2020-10-16 [1] CRAN (R 4.1.0)
#>  dotCall64              1.0-1     2021-02-11 [1] CRAN (R 4.1.0)
#>  dplyr                  1.0.5     2021-03-05 [1] CRAN (R 4.1.0)
#>  dqrng                  0.2.1     2019-05-17 [1] CRAN (R 4.1.0)
#>  DropletUtils           1.11.12   2021-02-21 [1] Bioconductor
#>  DT                     0.18      2021-04-14 [1] CRAN (R 4.1.0)
#>  edgeR                  3.33.3    2021-03-09 [1] Bioconductor
#>  ellipsis               0.3.1     2020-05-15 [1] CRAN (R 4.1.0)
#>  evaluate               0.14      2019-05-28 [1] CRAN (R 4.1.0)
#>  ExperimentHub          1.99.5    2021-04-16 [1] Bioconductor
#>  fansi                  0.4.2     2021-01-15 [1] CRAN (R 4.1.0)
#>  farver                 2.1.0     2021-02-28 [1] CRAN (R 4.1.0)
#>  fastmap                1.1.0     2021-01-25 [1] CRAN (R 4.1.0)
#>  fields                 11.6      2020-10-09 [1] CRAN (R 4.1.0)
#>  filelock               1.0.2     2018-10-05 [1] CRAN (R 4.1.0)
#>  foreach                1.5.1     2020-10-15 [1] CRAN (R 4.1.0)
#>  fs                     1.5.0     2020-07-31 [1] CRAN (R 4.1.0)
#>  generics               0.1.0     2020-10-31 [1] CRAN (R 4.1.0)
#>  GenomeInfoDb         * 1.27.11   2021-04-08 [1] Bioconductor
#>  GenomeInfoDbData       1.2.4     2020-11-03 [1] Bioconductor
#>  GenomicAlignments      1.27.2    2020-12-12 [1] Bioconductor
#>  GenomicRanges        * 1.43.4    2021-04-04 [1] Bioconductor
#>  ggbeeswarm             0.6.0     2017-08-07 [1] CRAN (R 4.1.0)
#>  ggplot2                3.3.3     2020-12-30 [1] CRAN (R 4.1.0)
#>  glue                   1.4.2     2020-08-27 [1] CRAN (R 4.1.0)
#>  golem                  0.3.1     2021-04-17 [1] CRAN (R 4.1.0)
```

```
#>  gridExtra                2.3       2017-09-09 [1] CRAN (R 4.1.0)
#>  gtable                   0.3.0     2019-03-25 [1] CRAN (R 4.1.0)
#>  HDF5Array                1.19.15   2021-04-25 [1] Bioconductor
#>  htmltools                0.5.1.1   2021-01-22 [1] CRAN (R 4.1.0)
#>  htmlwidgets              1.5.3     2020-12-10 [1] CRAN (R 4.1.0)
#>  httpuv                   1.6.0     2021-04-23 [1] CRAN (R 4.1.0)
#>  httr                     1.4.2     2020-07-20 [1] CRAN (R 4.1.0)
#>  interactiveDisplayBase   1.29.0    2020-10-27 [1] Bioconductor
#>  IRanges                * 2.25.10   2021-04-22 [1] Bioconductor
#>  irlba                    2.3.3     2019-02-05 [1] CRAN (R 4.1.0)
#>  iterators                1.0.13    2020-10-15 [1] CRAN (R 4.1.0)
#>  jquerylib                0.1.4     2021-04-26 [1] CRAN (R 4.1.0)
#>  jsonlite                 1.7.2     2020-12-09 [1] CRAN (R 4.1.0)
#>  KEGGREST                 1.31.1    2020-11-23 [1] Bioconductor
#>  knitr                    1.33      2021-04-24 [1] CRAN (R 4.1.0)
#>  labeling                 0.4.2     2020-10-20 [1] CRAN (R 4.1.0)
#>  later                    1.2.0     2021-04-23 [1] CRAN (R 4.1.0)
#>  lattice                  0.20-41   2020-04-02 [1] CRAN (R 4.1.0)
#>  lazyeval                 0.2.2     2019-03-15 [1] CRAN (R 4.1.0)
#>  lifecycle                1.0.0     2021-02-15 [1] CRAN (R 4.1.0)
#>  limma                    3.47.12   2021-03-28 [1] Bioconductor
#>  locfit                   1.5-9.4   2020-03-25 [1] CRAN (R 4.1.0)
#>  lubridate                1.7.10    2021-02-26 [1] CRAN (R 4.1.0)
#>  magick                   2.7.1     2021-03-20 [1] CRAN (R 4.1.0)
#>  magrittr                 2.0.1     2020-11-17 [1] CRAN (R 4.1.0)
#>  maps                     3.3.0     2018-04-03 [1] CRAN (R 4.1.0)
#>  Matrix                   1.3-2     2021-01-06 [1] CRAN (R 4.1.0)
#>  MatrixGenerics         * 1.3.1     2021-02-01 [1] Bioconductor
#>  matrixStats            * 0.58.0    2021-01-29 [1] CRAN (R 4.1.0)
#>  memoise                  2.0.0     2021-01-26 [1] CRAN (R 4.1.0)
#>  mime                     0.10      2021-02-13 [1] CRAN (R 4.1.0)
#>  munsell                  0.5.0     2018-06-12 [1] CRAN (R 4.1.0)
#>  pillar                   1.6.0     2021-04-13 [1] CRAN (R 4.1.0)
#>  pkgconfig                2.0.3     2021-03-31 [1] Github (gaborcsardi/pkgconfig@b81ae03)
#>  pkgload                  1.2.1     2021-04-06 [1] CRAN (R 4.1.0)
#>  plotly                   4.9.3     2021-01-10 [1] CRAN (R 4.1.0)
#>  plyr                     1.8.6     2020-03-03 [1] CRAN (R 4.1.0)
#>  png                      0.1-7     2013-12-03 [1] CRAN (R 4.1.0)
#>  Polychrome               1.2.6     2020-11-11 [1] CRAN (R 4.1.0)
#>  promises                 1.2.0.1   2021-02-11 [1] CRAN (R 4.1.0)
#>  pryr                   * 0.1.4     2018-02-18 [1] CRAN (R 4.1.0)
#>  purrr                    0.3.4     2020-04-17 [1] CRAN (R 4.1.0)
#>  R.methodsS3              1.8.1     2020-08-26 [1] CRAN (R 4.1.0)
#>  R.oo                     1.24.0    2020-08-26 [1] CRAN (R 4.1.0)
#>  R.utils                  2.10.1    2020-08-26 [1] CRAN (R 4.1.0)
#>  R6                       2.5.0     2020-10-28 [1] CRAN (R 4.1.0)
#>  rappdirs                 0.3.3     2021-01-31 [1] CRAN (R 4.1.0)
#>  RColorBrewer             1.1-2     2014-12-07 [1] CRAN (R 4.1.0)
#>  Rcpp                     1.0.6     2021-01-15 [1] CRAN (R 4.1.0)
#>  RCurl                    1.98-1.3  2021-03-16 [1] CRAN (R 4.1.0)
#>  RefManageR             * 1.3.0     2020-11-03 [1] Github (ropensci/RefManageR@ab8fe60)
```

```
#>  remotes                  2.3.0      2021-04-01 [1] CRAN (R 4.1.0)
#>  restfulr                 0.0.13     2017-08-06 [1] CRAN (R 4.1.0)
#>  rhdf5                    2.35.2     2021-02-18 [1] Bioconductor
#>  rhdf5filters             1.3.4      2021-02-17 [1] Bioconductor
#>  Rhdf5lib                 1.13.4     2021-02-18 [1] Bioconductor
#>  rjson                    0.2.20     2018-06-08 [1] CRAN (R 4.1.0)
#>  rlang                    0.4.10     2020-12-30 [1] CRAN (R 4.1.0)
#>  rmarkdown                2.7        2021-02-19 [1] CRAN (R 4.1.0)
#>  roxygen2                 7.1.1      2020-06-27 [1] CRAN (R 4.1.0)
#>  rprojroot                2.0.2      2020-11-15 [1] CRAN (R 4.1.0)
#>  Rsamtools                2.7.2      2021-04-10 [1] Bioconductor
#>  RSQLite                  2.2.7      2021-04-22 [1] CRAN (R 4.1.0)
#>  rstudioapi               0.13       2020-11-12 [1] CRAN (R 4.1.0)
#>  rsvd                     1.0.5      2021-04-16 [1] CRAN (R 4.1.0)
#>  rtracklayer            * 1.51.5     2021-03-09 [1] Bioconductor
#>  S4Vectors              * 0.29.17    2021-04-23 [1] Bioconductor
#>  sass                     0.3.1.9000 2021-02-12 [1] Github (rstudio/sass@2526470)
#>  ScaledMatrix             0.99.2     2021-01-14 [1] Bioconductor
#>  scales                   1.1.1      2020-05-11 [1] CRAN (R 4.1.0)
#>  scater                   1.19.11    2021-02-26 [1] Bioconductor
#>  scatterplot3d            0.3-41     2018-03-14 [1] CRAN (R 4.1.0)
#>  scuttle                  1.1.18     2021-02-26 [1] Bioconductor
#>  sessioninfo            * 1.1.1      2018-11-05 [1] CRAN (R 4.1.0)
#>  shiny                    1.6.0      2021-01-25 [1] CRAN (R 4.1.0)
#>  shinyWidgets             0.6.0      2021-03-15 [1] CRAN (R 4.1.0)
#>  SingleCellExperiment   * 1.13.14    2021-04-07 [1] Bioconductor
#>  spam                     2.6-0      2020-12-14 [1] CRAN (R 4.1.0)
#>  sparseMatrixStats        1.3.7      2021-03-29 [1] Bioconductor
#>  SpatialExperiment      * 1.1.5      2021-04-28 [1] Github (drighelli/SpatialExperiment@17e5176)
#>  spatialLIBD            * 1.3.15     2021-04-29 [1] Bioconductor
#>  stringi                  1.5.3      2020-09-09 [1] CRAN (R 4.1.0)
#>  stringr                  1.4.0      2019-02-10 [1] CRAN (R 4.1.0)
#>  SummarizedExperiment   * 1.21.3     2021-04-07 [1] Bioconductor
#>  testthat                 3.0.2      2021-02-14 [1] CRAN (R 4.1.0)
#>  tibble                   3.1.1      2021-04-18 [1] CRAN (R 4.1.0)
#>  tidyr                    1.1.3      2021-03-03 [1] CRAN (R 4.1.0)
#>  tidyselect               1.1.0      2020-05-11 [1] CRAN (R 4.1.0)
#>  usethis                  2.0.1      2021-02-10 [1] CRAN (R 4.1.0)
#>  utf8                     1.2.1      2021-03-12 [1] CRAN (R 4.1.0)
#>  vctrs                    0.3.7      2021-03-29 [1] CRAN (R 4.1.0)
#>  vipor                    0.4.5      2017-03-22 [1] CRAN (R 4.1.0)
#>  viridis                  0.6.0      2021-04-15 [1] CRAN (R 4.1.0)
#>  viridisLite              0.4.0      2021-04-13 [1] CRAN (R 4.1.0)
#>  withr                    2.4.2      2021-04-18 [1] CRAN (R 4.1.0)
#>  xfun                     0.22       2021-03-11 [1] CRAN (R 4.1.0)
#>  XML                      3.99-0.6   2021-03-16 [1] CRAN (R 4.1.0)
#>  xml2                     1.3.2      2020-04-23 [1] CRAN (R 4.1.0)
#>  xtable                   1.8-4      2019-04-21 [1] CRAN (R 4.1.0)
#>  XVector                  0.31.1     2020-12-12 [1] Bioconductor
#>  yaml                     2.2.1      2020-02-01 [1] CRAN (R 4.1.0)
#>  zlibbioc                 1.37.0     2020-10-27 [1] Bioconductor
```

```
#>
#> [1] /Library/Frameworks/R.framework/Versions/4.1/Resources/library
```
